# Valorisation of spent cultivated meat media for recombinant FGF2 production in GRAS *Lactococcus lactis*

**DOI:** 10.1101/2024.08.01.606190

**Authors:** Juliana Rizal, Prashant Mainali, Jun Ping Quek, Tan Lee Ling, Jiawu Bi, Alson Jianchen Chan, Azra Anwar Gaffoor, Lamony Jian Ming Chew, Shigeki Sugii, Say Kong Ng, Dave Siak-Wei Ow, Fong Tian Wong

## Abstract

Innovative strategies for sustainable utilization of waste resources are imperative in the pursuit of a circular economy. Recently, the idea of utilizing mammalian spent media as a valuable resource is gaining traction, offering significant opportunities for innovative uses as a food-grade feedstock for microbial fermentation, especially in the production of alternative proteins for research and food purposes.

In this study, we aim to repurpose spent mammalian culture media for production of valuable proteins. Growth factors (GFs) are a family of high-value proteins that naturally stimulate cell proliferation or differentiation. More importantly, these factors also present significant costs for cell culture. Here, we successfully demonstrate the use of spent mammalian culture media for the recombinant production of fibroblast growth factor 2 (FGF2-G3) in *Lactococcus lactis*. Bioreactor fermentation at a 1 L scale confirmed purified yields of 2.6 mg/L of recombinant FGF2-G3 using spent media. Further functional testing indicated that the recombinant FGF2-G3 can promote cell proliferation on an *Anguilla japonica* (Japanese eel) pre-adipocytic cell line, suggesting its potential for cultivated meat production. Based on the preliminary results of this study, our calculations indicate that fermenting 1 L spent mammalian waste could yield enough growth factors to efficiently grow approximately 52 L of cultivated meat through fermentation. This prediction emphasizes the potential of waste valorisation to sustainably produce protein, thus contributing to environmental preservation and economic viability.

**Highlights:**

1. Spent mammalian culture media can be repurposed for microbial fermentation.
2. GRAS *L. lactis* was used to express FGF2-G3 in an optimized spent media formulation.
3. Functional FGF2-G3 expression was demonstrated in controlled 1-L bioreactor runs.

## Introduction

With increasing focus and bio-production via mammalian cultured systems for therapeutics to food, the concept of defined mammalian spent media emerges as a relatively unexplored yet rapidly expanding resource. This presents immense opportunities for innovation as a novel food-grade feedstock for microbial fermentation, particularly for precise fermentation aimed at alternative protein production towards research or food applications. Mammalian culture media are usually characterized by high quality standards (GMP-grade) and contain valuable nutrients, including amino acids and vitamins (Haraguchi et al., 2022). Furthermore, levels of pyruvate, vitamins, and inorganic salts do not significantly change after culturing (Haraguchi et al., 2022). In this study, our target is to upcycle spent mammalian culture media for the production of high value proteins. Growth factors (GFs) are naturally occurring substances that activate cells for proliferation or differentiation. The presence of mitogenic GFs is an essential prerequisite for cell culture medium, playing a crucial role in diverse applications within academia and the bioindustry. This significance extends to emerging fields like tissue engineering and cellular agriculture. Notable examples include epidermal growth factor (EGF) and fibroblast growth factor (FGF). However, the production of growth factors for cell culture medium is expensive. A pivotal component in culture media, these growth factors are vital for the maintenance and differentiation of cultured cells. Depending on the quality grade, the estimated cost of 1 gram of FGF-2 is $2 million, and TGF-beta is $80 million, constituting at least 96% of the cost of a commercial serum-free cell culture medium. A technical analysis conducted by The Good Food Institute in 2020 estimate the cost of the commercial serum-free media (Liquid Essential 8 Media) for cultivated meat (CM) in a hypothetical 20-tonne batch run to cost around USD$377 USD per litre, with growth factors accounting for 99% of the cost, based on state of the technology in 2020 (Specht, 2020). Two recent works have also demonstrated the use of spent media for cultivation of microalgae and *Escherichia coli* (Haraguchi et al., 2022 and Lynch & O’Connell, 2022).

FGF2 is a crucial member of the fibroblast growth factor family, that are involved in tissue development, differentiation and maintenance of cultured cells (Figure 1A, Enriquez-Ochoa et al., 2020). Through binding to specific receptors and inducing downstream signaling pathways, FGF2 regulates cellular behavior essential for proper tissue development and repair mechanisms. FGF2’s applications extend to biomedical research, simplifying stem cell culture protocols and supporting critical functions in stem cell research and regenerative medicine. FGF2 is also one of the typical growth factors that acts as a supplement for hESC culture media which helps to promote self-renewal of cultured cells (Eselleova, 2009). Here, we focus on FGF2-G3, which is a modified and stabilized version of FGF2 that possessed nine mutations (Dvorak et al, 2018). This variant has been shown to be more effective than previously identified variant, FGF2-K128N. In a short-term growth assay, 5 ng/mL of FGF2-G3 demonstrated similar growth-promoting effects as 100 ng/mL of FGF2-K128N, indicating a significant increase in potency and efficiency. This enhancement is particularly important in applications requiring precise and effective growth factor concentrations.

**Figure 1.**
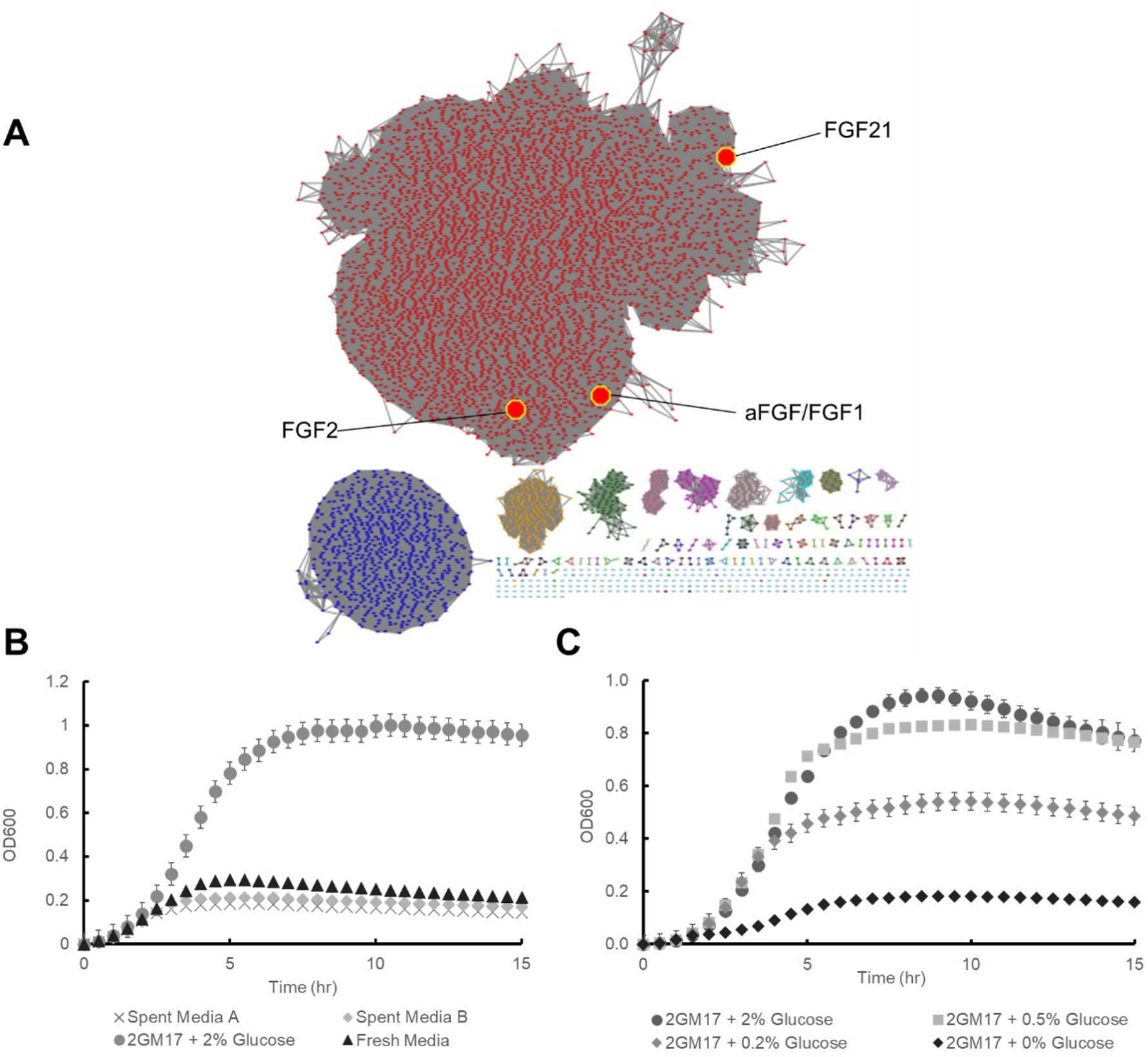
**(A)**. Sequence Similarity Network (SSN) of the Fibroblast Growth Factor (FGF) Family. Closely related growth factors are grouped into the same clusters and colour-coded in the same colour. Within each cluster, distance between each node (protein member) and the number of edges (neighboring connections) are indicators of how close the proteins are to each other. aFGF: acidic fibroblast growth factor. FGF1, FGF2, FGF21 are fibroblast growth factors 1, 2 and 21 respectively. **(B)** Growth curves of *L. lactis* NZ9000 at 30ºC. Spent media A and media B are spent mammalian media from two different cultivation fermentation, fresh media is the in-house formulated chemically defined serum-free, protein-free growth media. These were performed in biological triplicates at 30°C with standard deviations as error bars. **(C)** Growth curves of *L. lactis* NZ9000 at 30ºC in the presence of different glucose concentrations (2%, 0.5%, 0.2% and 0%) in 2GM17 media. These were conducted in biological triplicates at 30ºC with standard deviations as error bars.

To simplify food grade operations, our organism of choice is *Lactococcus lactis*, a well-known lactic acid bacteria (LAB). It is a Gram-positive bacteria which does not produce endotoxins and unwanted glycosylation of proteins (Singh et al., 2018). It holds the distinction of being recognized as Generally Regarded as Safe (GRAS). This bacterial species is extensively utilized as starter cultures in fermentation processes and serves a crucial role as probiotics (Lim et al., 2017). Moreover, its industrial applications extend to fermentations, where it contributes to enhancing the taste and texture of food and feed (Mokoena et al., 2021). In the context of medical applications, LAB is also an ideal and inexpensive live vector for *in-situ* delivery of recombinant therapeutic proteins (del Rio, 2019). It is interesting to note that current examples of growth factor expression in LAB have mainly been for *in-situ* delivery (FGF21: Cao, W-Y., et al.; EGF: Huynh, Evanna and Julang Li). Encouragingly, recent work on the secretory expression of gallus epidermal growth factor (gEGF) managed to demonstrate activity. Based on supernatant from these fermentations, it was sufficient to stimulate the proliferation of chicken embryo fibroblast cells (Zhou, Y. 2021).

In this study, we are presenting the first instance of recombinant FGF2-G3 expression in *L. lactis*. Although FGF21 production has been demonstrated previously, this demonstration of FGF2-G3 production, another distinct member of the FGF family (Figure 1A), expands the potential of *L. lactis* for FGF expression. Additionally, we have successfully demonstrated the efficient repurposing of an optimized spent serum-free mammalian culture media for the recombinant production of functional FGF2-G3 in GRAS *L. lactis*. Moreover, we have scaled up this process for production in bioreactors. These initial findings suggest the potential for precise fermentation of GRAS organisms using spent mammalian waste as part of strategies to further reduce the overall costs of cultivated meat production.

## Materials and methods

### Bacteria strains and protein expression plasmids

*L. lactis* NZ9000 and pNZ8148 vector (Boca Scientific, USA) were used for this investigation. FGF2-G3 gene (Dvorak et al., 2018) was codon optimized and placed under nisin-inducible promoter, Pnis, in plasmid pNZ8148 (Boca Scientific, USA). FGF2-G3 has been His-tagged to facilitate protein analysis and purification.

### Measurement of growth rates

Growth was monitored over a 15-hour incubation period at 30°C, where OD600 readings were taken at 30 minutes intervals using a Varioskan™ LUX multimode microplate reader. Biological triplicates were performed for growth measurements.

### FGF2 expression and purification

250 µL overnight culture (2.5%) was used to inoculate 10 mL media. The cultures were grown at 30°C until OD600 = 0.5-0.6, before induction with nisin at a final concentration of 50 ng/mL. Next, the cultures were incubated overnight. Cells were harvested using a microcentrifuge at 8,000 x g at 4°C for 10 minutes before resuspending the cell pellet in 500 µL 50 mM sodium phosphate buffer (pH = 7.4), 300 mM sodium chloride, 10 mM imidazole and 0.03 % Triton X-100. Cells were lysed with sonication. The lysed cells were then centrifuged at maximum speed for 10 minutes at 4°C. The resulting supernatant was collected and loaded onto 100 µL Ni-NTA resin. This was washed with 600 µL 50 mM sodium phosphate (pH=7), 300 mM sodium chloride, 50 mM imidazole. 100 µL 50 mM sodium phosphate (pH=7), 300 mM sodium chloride, 500 mM imidazole was used for elution. Eluted proteins were then quantified with SDS-PAGE. Biological triplicates were performed.

### Bioreactor fermentation and purification

Fermentation was conducted in a 1 L Biostat A bioreactor (Sartorius Stedim, Germany). The experiment was carried out with a working volume of 500 mL. The pH was maintained at 7 using a PID controller, which adjusted the pH with 14 % NH4OH and 1 M H2SO4 as needed. Dissolved oxygen was kept at 0% saturation by sparging nitrogen gas at the beginning of the experiment. A constant stirrer speed of 150 rpm and a temperature of 30°C were maintained throughout the run. The initial cell density at the start of the experiment was 0.15 OD. When the cell density reached approximately 0.5 OD, the cells were induced with 50 ng/ml of nisin. At each sampling point, 11 mL of cell culture was withdrawn: 1 mL was used to measure the OD, and the remaining 10 ml was centrifuged to pellet the cells for FGF2-G3 protein extraction. At final timepoint, 25 mL of cell volume was pelleted and purified through His-tag affinity chromatography and buffer exchanged into PBS in duplicates. Protein yield of FGF2–G3 was quantified using Bradford assays.

### Cell line and cell cultivation

Chicken fibroblast cell, UMNSAH/DF-1 (ATCC® CRL12203™) were obtained from the American Type Culture Collection (ATCC) and adapted to suspension culture in an in-house proprietary Dulbecco’s modified Eagle’s medium (DMEM)/F12-based protein-free chemically defined medium. Cells were cultured in disposable Erlenmeyer flasks (Corning, USA) in a humidified shaking incubator (Kuhner, Switzerland) at 39°C, 5% CO2, and 110 rpm. For routine cultivation, cells were passaged every 3–4 days by seeding at 5 × 10^5^ cells/ml. Viable cell densities and viabilities were determined using Vi-CELL XR Cell Viability Analyzer (Beckman Coulter, USA), according to manufacturer’s instructions. Spent media was collected every 3– 4 days, filtered via 0.22mM vacuum filtration systems (Corning, USA) and analyse by BioProfile 100 Plus (Nova Biomedical, USA).

*Anguilla japonica* pre-adipocytic cell line, Aj1C-2x was isolated with the method as described by Sugii et al., 2011 with modifications. Briefly, intramuscular fat of *Anguilla japonica* was digested with equal volume (v/w) of HBSS containing 1 mg/ml of collagenase type IV (Worthington Biochemical Corporation, NJ, USA) and 5 mM calcium chloride in 37°C for 1 hour. After several passages, the stromal vascular fraction was sorted into 96-well plate at 1 cell per well using BD FACSAria II Cell Sorter (BD Biosciences, USA). The cells were spontaneously immortalized by passaging for more than 30 passages. The cells line was adapted and cultured in Dulbecco’s modified Eagle’s medium (DMEM)/F12 media (Thermo Fisher Scientific, USA) supplemented with 2.5% reduced fetal bovine serum and 10 ng/mL of FGF2 in a humidified incubator controlled at 27°C and supplied with 5% CO2. Viable cell densities and viabilities were determined using Vi-CELL XR Cell Viability Analyzer (Beckman Coulter, USA), according to manufacturer’s instructions. All procedures involving live animal handling were performed as approved by the Institutional Animal Care and Use Committee (IACUC) of A*STAR, Singapore (IACUC no. #201570).

### Growth factor bioactivity assay

Aj1C-2x cells were seeded in a 96-well plate at a seeding density of 2e4 cells/well, DMEM/12 supplemented with 2.5% FBS and varying concentration of commercial heat stable FGF2 and recombinant FGF2-G3 and cultured for 3 days. Commercial heat stable FGF2 (PHG0368, Thermo Fisher Scientific) was used as positive control. After 3 days of incubation, CyQuant XTT cell viability assay (X12223, Thermo Fisher Scientific) was performed according to the manufacturer’s instructions. To determine the specific absorbance reading of each sample, the absorbance reading of the media control was subtracted from the absorbance reading of each sample. Subsequently, to calculate the relative absorbance fold change, the specific absorbance reading of each sample were normalized against the specific absorbance reading of the cells cultivated in basal media with 2.5% FBS for Aj1C-2x cells.

## Results

### Chemically defined serum free media is unable to support LAB growth

DF-1 cells (ATCC CRL-12203™) were suspension-adapted and cultured in in-house formulated chemically defined serum-free, protein-free growth media. After 3 - 4 days of cell culture, the spent media containing depleted nutrients and accumulated waste metabolites were harvested. To examine the content of spent media waste, we analysed 7 batches of >100 mL fermentation waste using Bioprofile 100 plus Analyser (Table 1). Notably, compared to fresh media, there is a drop in glucose, increase in lactate and ammonia waste concentration. To first examine if these media was sufficient to support growth of *L. lactis*, growth rates of *L. lactis* strain, NZ9000, was measured in (1) rich microbial media (2GM17 + 2% glucose; Lim et al., 2017), (2) fresh mammalian media and (3) spent serum-free mammalian media (spent media A and B) (Figure 1B). As glucose is important for *L. lactis* cultivation, we compared the growth of *L. lactis* under various glucose concentrations using 2GM17 media. These concentrations ranged from 0 to 2%. Our findings indicated that 0.2% glucose (similar to that of spent media, Table 1) should be sufficient to support growth. Although the chemically defined mammalian media is still carbon-rich and contains approximately 0.2 g/L glucose, we found that it was unable to support growth of *L. lactis*, indicating additional nutritional requirements.

**Table 1.** Metabolite profile of fresh media and spent media. Metabolite concentrations were quantified using Bioprofile 100 plus Analyser.

| Components (units) | Fresh Media | Spent media (n=7) |  |
| --- | --- | --- | --- |
|  |  | Average | stdev |
| pH | 7.34 | 6.78 | 0.12 |
| Glutamine (mM) | 3.04 | 1.78 | 0.21 |
| Glutamate (mM) | 0.56 | 0.49 | 0.12 |
| Glucose (g/l) | 3.39 | 1.95 | 0.17 |
| Lactate (g/l) | 0 | 1.53 | 0.13 |
| NH<sub>4</sub><sup>+</sup> (mM) | 0.22 | 0.92 | 0.14 |
| Na<sup>+</sup> (mM) | 142 | 143 | 7 |
| K<sup>+</sup> (mM) | 4 | 4 | 0.2 |

### Production of FGF2-G3 in optimized spent media formulation

Subsequently to express FGF2-G3 in spent mammalian media, we added a proprietary mixture of non-sugar supplements to account for essential nutrients lacking in spent media. Under these conditions, growth of *L. lactis* was restored. Recombinant production of FGF2-G3 was also possible under these conditions; moreover, protein yields were comparable to 2GM17 + 2% glucose (Figure 2A, B).

**Figure 2.**
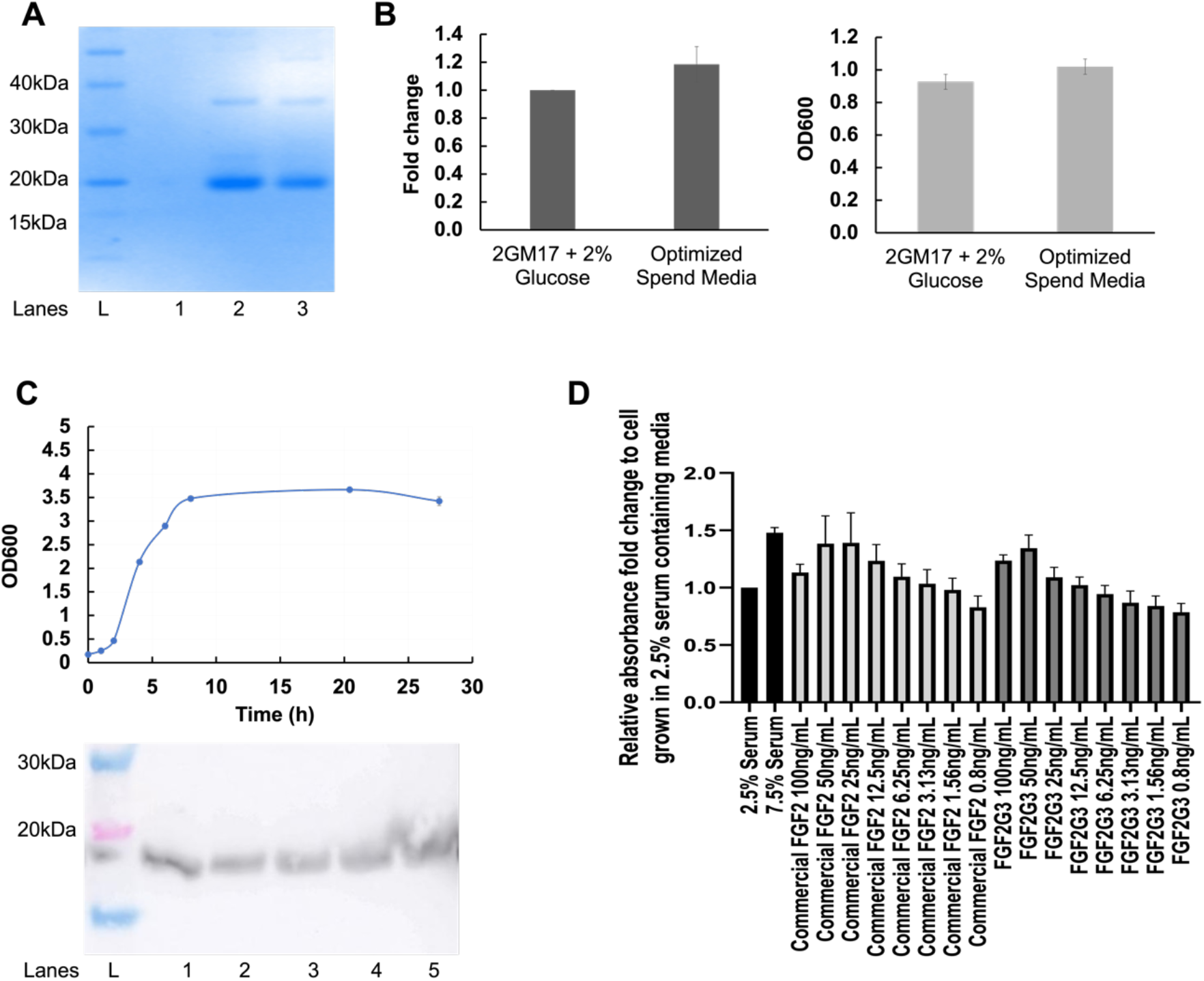
**(A)** Production of recombinant purified FGF2-G3 from *L. lactis* fermented in various media, visualized by SDS-PAGE. Lane 1 is spent media (no additives), Lane 2 is optimized spent media (spent media with additives) and Lane 3 is control rich media; 2GM17+2% glucose. Proteins were purified from 10 mL-scale cultures. These were conducted in biological triplicates at 30ºC with standard deviations as error bars. **(B)** Average fold change of protein yields and final OD600 of FGF2-G3 cultured in 2GM17+2% glucose and optimized spent media. Fold change was calculated by densitometry via ImageJ. Final OD600 was measured at 15 hour. **(C)** Production of recombinant FGF2-G3 in optimized spent media in 1 L Biostat A bioreactor (Sartorius Stedim, Germany) – growth curve (OD600) and production of recombinant FGF2-G3 in 1 L Biostat A bioreactor, visualized by Western Blot with anti-FGF2-G3 staining. Lane 1-5 are samples from 4, 6, 8, 20 and 27 hours post-induction respectively. **(D)** Cell viability assay was performed on Aj1C-2x cells, seeded in a 96-well plate at a seeding density of 2E4 cells/well, with varying concentration of commercial heat stable FGF2 (Thermo Fisher Scientific) and purified recombinant FGF2-G3. Absorbance readings are normalized to cells cultivated in DMEM/F12 media supplemented with 2.5% FBS. Data plotted as mean ± standard deviation (SD) using Graphpad Prism v8.0. Error bars represent the SD calculated for the replicates.

After successfully demonstrating the production of recombinant FGF2-G3 in optimized spent media with benchtop cultures, we proceeded with bioreactor fermentation. The goal of the 1 L bioreactor experiment was to verify if the optimized spent media formulation for recombinant protein production *L. lactis* was consistent at larger volumes and cell densities. The main difference between small-scale and bioreactor-scale experiments is the controlled environment for cell growth, pH (maintained at pH 7), and the level of dissolved oxygen (maintained at 0% saturation). It is anticipated that this controlled environment will lead to a higher final cell density at the end of fermentation.

During the fermentation with optimized spent media, the culture was allowed sufficient time to adjust to the less nutrient-rich media before induction (Figure 2C). The experiment began at an optical density (OD) of 0.15, and induction took place when the cells reached an OD of 0.5. After induction, the cells entered the stationary phase approximately 7 hours later. Majority of protein expression occurred within the first 2 hours post-induction, as shown by the Western blot results in Figure 2C, where the highest band intensity was observed at the 4-hour time point. At later time points, the band intensity either remained the same or decreased. The final purified FGF2-G3 concentration at the end of the 27.5-hour fermentation was 2.6 mg/L (± 0.14 mg/L).

In this study, FGF2-G3 was purified using a single affinity pull-down step. This takes advantage of the fact that *L. lactis* does not produce LPS and is a food-grade bacteria. Biological functional testing of the purified FGF2-G3 was conducted using the Unagi (Japanese eel) pre-adipocytic cell line, Aj1C-2x, with concentrations ranging from 0.08 to 100 ng/mL. Our observations revealed that both the commercial heat-stable FGF2 and the recombinant FGF2-G3 were able to promote cell proliferation better than cell growth in 2.5% fetal bovine serum (FBS) when used at a minimum concentration of 12.5 ng/mL and 25 ng/mL, respectively (Figure 2D). Based on a requirement of 50 ng/mL for our recombinant FGF2-G3, this would also mean that recombinant proteins fermented from 1 L fermentation of spent mammalian waste could provide sufficient growth factors for efficient cell proliferation of approximately 52 L of cultivated meat fermentation.

## Conclusion

Given that one milligram of commercial FGF2-G3 costs around 1,200 USD^14^, utilizing spent serum-free cultivated meat media to produce high-value growth factors at milligram levels using *L. lactis* can significantly reduce the cost of the final media and strongly supports a sustainable circular economy. Furthermore, our bioreactor fermentation has shown that the production and purification of FGF2-G3 in spent mammalian media is scalable and can yield FGF2-G3 that is commercially lucrative and relevant. While we have demonstrated the first example of recombinant protein production from GRAS *L. lactis* using spent mammalian media, further fermentation optimization, including control of nisin, induction time, and temperature, can be used to improve yields and productivity.

## Acknowledgements

NA

## Author contributions

**Juliana Rizal:** Methodology, Investigation, Data curation, Visualization, Writing-original draft, Writing-review and editing. **Prashant Mainali:** Methodology, Investigation, Data curation, Visualization, Writing-original draft, Writing-review and editing. **Lee Ling Tan:** Methodology **Jiawu Bi:** Visualization **Alson Jianchen Chan:** Methodology **Jun Ping Quek:** Methodology, Investigation, Data curation, Writing-original draft, Writing - review and editing **Azra Anwar Gaffoor:** Methodology **Lamony Jian Ming Chew:** Methodology, Data curation, Writing-original draft. **Shigeki Sugii:** Supervision, Writing-review and editing. **Say Kong Ng:** Conceptualization, funding acquisition, Writing-review and editing. **Dave Siak-Wei Ow:** Conceptualization, Funding acquisition, Supervision, Writing-review and editing. **Fong Tian Wong:** Conceptualization, Funding acquisition, Supervision, Writing-original draft, Writing-review and editing.

## Data availability

- Data will be made available on request.

## References

[1] Cao, W.-Y., Dong, M., Hu, Z.-Y., Wu, J., Li, Y.-C., & Xu, H.-D. (2020). Recombinant Lactococcus lactis NZ3900 expressing bioactive human FGF21 reduced body weight of Db/Db mice through the activity of brown adipose tissue. Beneficial Microbes, 11(1), 67–78. 10.3920/BM2019.0093

[2] del Rio, B., Redruello, B., Fernandez, M., Martin, M. C., Ladero, V., & Alvarez, M. A. (2019). Lactic Acid Bacteria as a Live Delivery System for the in situ Production of Nanobodies in the Human Gastrointestinal Tract. Frontiers in Microbiology, 9. 10.3389/fmicb.2018.03179

[3] Dvorak, P., Bednar, D., Vanacek, P., Balek, L., Eiselleova, L., Stepankova, V., Sebestova, E., Kunova Bosakova, M., Konecna, Z., Mazurenko, S., Kunka, A., Vanova, T., Zoufalova, K., Chaloupkova, R., Brezovsky, J., Krejci, P., Prokop, Z., Dvorak, P., & Damborsky, J. (2018). Computer-assisted engineering of hyperstable fibroblast growth factor 2. Biotechnology and Bioengineering, 115(4), 850–862. 10.1002/bit.26531

[4] Eiselleova, L., Matulka, K., Kriz, V., Kunova, M., Schmidtova, Z., Neradil, J., Tichy, B., Dvorakova, D., Pospisilova, S., Hampl, A., & Dvorak, P. (2009). A complex role for FGF-2 in self-renewal, survival, and adhesion of human embryonic stem cells. Stem Cells (Dayton, Ohio), 27(8), 1847–1857. 10.1002/stem.128

[5] Enriquez-Ochoa, D., Robles-Ovalle, P., Mayolo-Deloisa, K., & Brunck, M. E. G. (2020). Immobilization of Growth Factors for Cell Therapy Manufacturing. Frontiers in Bioengineering and Biotechnology, 8. 10.3389/fbioe.2020.00620

[6] Haraguchi, Y., Okamoto, Y., & Shimizu, T. (2022). A circular cell culture system using microalgae and mammalian myoblasts for the production of sustainable cultured meat. Archives of Microbiology, 204(10), 615. 10.1007/s00203-022-03234-9

[7] Lim, P. Y., Tan, L. L., Ow, D. S.-W., & Wong, F. T. (2017). A propeptide toolbox for secretion optimization of Flavobacterium meningosepticum endopeptidase in Lactococcus lactis. Microbial Cell Factories, 16(1), 221. 10.1186/s12934-017-0836-0

[8] Lynch, C. D., & O’Connell, D. J. (2022). Conversion of mammalian cell culture media waste to microbial fermentation feed efficiently supports production of recombinant protein by Escherichia coli. PLOS ONE, 17(5), e0266921. 10.1371/journal.pone.0266921

[9] Mokoena, M. P., Omatola, C. A., & Olaniran, A. O. (2021). Applications of Lactic Acid Bacteria and Their Bacteriocins against Food Spoilage Microorganisms and Foodborne Pathogens. Molecules, 26(22), 7055. 10.3390/molecules26227055

[10] Singh, S. K., Tiendrebeogo, R. W., Chourasia, B. K., Kana, I. H., Singh, S., & Theisen, M. (2018). Lactococcus lactis provides an efficient platform for production of disulfide-rich recombinant proteins from Plasmodium falciparum. Microbial Cell Factories, 17(1), 55. 10.1186/s12934-018-0902-2

[11] Specht, L. (2020). An analysis of culture medium costs and production volumes for cultivated meat.

[12] Sugii, S., Kida, Y., Berggren, W. T., & Evans, R. M. (2011). Feeder-dependent and feeder-independent iPS cell derivation from human and mouse adipose stem cells. Nature Protocols, 6(3), 346–358. 10.1038/nprot.2010.199

[13] Zhou, Y., Chen, P., Shi, S., Li, X., Shi, D., Zhou, Z., Li, Z., & Xiao, Y. (2021). Expression of Gallus Epidermal Growth Factor (gEGF) with Food-Grade Lactococcus lactis Expression System and Its Biological Effects on Broiler Chickens. Biomolecules, 11(1), 103. 10.3390/biom11010103

[14] ThermoFisher Scientific. Human Heat Stable bFGF Recombinant Protein, expressed in E. coli. Retrieved July 24, 2024, from https://www.thermofisher.com/proteins/product/Human-Heat-Stable-bFGF-Recombinant-Protein/PHG0369

